# The Effects of Oral Contraceptives on Grip Strength and Neuromuscular Activation During Short-Term Immobilization and Rehabilitation of the Wrist/Hand

**DOI:** 10.1101/2025.07.01.662432

**Authors:** Matt S. Stock, Hannah M. Bauerlein, Maya F. Edwards, Kelsey E. Stambaugh, Emily J. Parsowith, Joshua C. Carr, Abbie E. Smith-Ryan, Randi M. Richardson

## Abstract

Females often experience greater weakness following immobilization compared to males. Hormonal fluctuations from the menstrual cycle or oral contraceptive (OC) use may contribute to sex differences and response variation. We examined differences in peak and rapid force and neuromuscular activation among females using monophasic OC and females not using OC following immobilization and rehabilitation. To examine potential sex differences, a male control group was included. Ten males, 10 OC females, and 10 non-OC females (mean ± SD age = 23 ± 3 years) immobilized their left wrist/hand with a brace for one week, followed by ≥ one week of rehabilitation. Participants completed grip tests to assess peak force and the rate of force development (RFD) before and after immobilization and post-rehabilitation, with electromyographic signals recorded from the extensor carpi radialis brevis (ECBR) and flexor digitorum superficialis (FDS). Grip force declined post-immobilization: males = –17.2 ± 10.3%, non-OC = –22.3 ± 24.7%, OC = –20.7 ± 14.8%. No significant time × group interactions were observed for any dependent variables (p > 0.05, η^2^_p_ ≤ 0.084). Time effects showed recovery post-rehab. RFD, particularly at 200 ms, declined posttest and rebounded post-rehab. ECBR excitation increased post-rehab; FDS trended upward. Only 5 participants required > one week of rehabilitation (2 males, 2 non-OC, 1 OC). We conclude that males and females exhibit similar declines and recovery in grip force after one week of wrist/hand immobilization, regardless of OC use. OC use does not appear to affect outcomes in female patients undergoing musculoskeletal rehabilitation.

**NEW & NOTEWORTHY:** Previous studies report greater strength losses in females than males after immobilization, yet the underlying mechanisms remain unclear. This is the first study to examine whether monophasic OC use influences neuromuscular responses to disuse and recovery. We found no significant differences in the loss or recovery of peak force, RFD, or sEMG excitation between OC users, non-users, and males. Short-term immobilization and rehabilitation outcomes are not meaningfully affected by OC use in healthy, young females.

## INTRODUCTION

The losses of muscle strength and size are clinically significant consequences of disuse, frequently observed after orthopedic injury, surgery, or prolonged hospitalization (1). Strength losses often exceed the degree of atrophy and can occur rapidly (2), driven by neuromuscular impairments such as reduced voluntary activation (3), altered motor unit recruitment (4), and diminished brain activity (5). Muscle atrophy, in contrast, tends to be slower and is primarily governed by increases in protein degradation (6) and increased activation of proteolytic pathways (7). Together, these neural and muscular adaptations contribute to substantial functional impairments that delay recovery and increase the risk of long-term disability. These issues present major challenges for healthcare professionals—especially when patients fail to return to their functional baseline, placing greater demands on rehabilitation services and long-term care. A better understanding of the factors contributing to poor recovery may support the development of more targeted exercise and rehabilitation strategies.

Prior studies indicate that females experience greater declines in muscle function during joint immobilization compared to males, despite smaller sex differences in muscle atrophy (8–12). The reason(s) for sex-specific strength losses following immobilization have not been thoroughly studied, but may include greater declines in neuromuscular function among females (10). Even less is known about sex-specific recovery following disuse, with only two studies to date directly examining differences between males and females in recovery following joint immobilization (12,13). Clark et al. (13) examined wrist flexion strength after three weeks of wrist/hand immobilization in 5 males and 5 females. Although strength losses were similar between sexes during immobilization, only males recovered to baseline after one week, while females showed minimal improvement. Muscle size was not assessed; however, unchanged central activation suggested the observed differences were peripheral in origin. More recently, Girts et al. (12) reported a 10.8% decrease in quadriceps peak torque in males and a 15.2% decrease in females following one week of knee immobilization. Participants resumed resistance training twice weekly until strength returned to baseline. While not statistically significant, females required more retraining sessions (mean = 2.91; median = 2) than males (mean = 2.13; median = 1) to restore peak torque. Notably, muscle atrophy persisted at the end of the study, underscoring a dissociation between muscle size and strength during recovery. Together, these findings suggest that females exhibit greater impairments during immobilization and slower recovery with or without rehabilitation compared to males, although the underlying mechanisms remain poorly understood.

Fluctuations in ovarian hormones across the menstrual cycle may influence neuromuscular responses to disuse and recovery, potentially contributing to poorer outcomes in females. Estrogen generally exerts excitatory effects on the central nervous system, enhancing glutamatergic signaling and suppressing GABAergic inhibition (14), whereas progesterone has opposing, inhibitory actions that may reduce motor unit firing rates and voluntary activation (15). These hormonal shifts are particularly relevant during the luteal phase, when progesterone is elevated and has been shown to suppress low-threshold motor unit firing rates, alter recruitment strategies, and increase intracortical inhibition (15,16). While cycle-dependent hormonal fluctuations may alter neuromuscular responses via central or peripheral mechanisms, this has never been evaluated in the disuse and rehabilitation literature. It is possible that previously reported sex differences were influenced by a failure to account for the role of the menstrual cycle in immobilization-induced weakness and recovery.

Similar to gaps in knowledge pertaining to the role of the menstrual cycle, no previous immobilization studies have accounted for the use of hormonal contraceptives. Over 60% of females in the United States use some form of hormonal contraceptive (17), making this a highly relevant factor to consider in studies involving neuromuscular adaptations. Among these, oral contraceptive (OC) use is one of the most common methods—reported by approximately 14% of U.S. females aged 15–49 as of 2017–2019, according to CDC data (18). Despite its prevalence, the physiological effects of OC use on skeletal muscle and neuromuscular adaptations are not well understood. OCs alter circulating levels of endogenous hormones such as estradiol and progesterone, which are known to exert a variety of physiological effects. These hormonal shifts may have implications for how females respond to both disuse and recovery. However, the effects of OC use on neuromuscular adaptation to resistance training remain unclear (19), with some studies suggesting potential blunting of responses (20), while others report favorable outcomes (21,22). These limited, mixed findings highlight a critical gap in our understanding of how OCs influence neuromuscular plasticity, particularly in the context of immobilization and recovery.

Understanding the impact of exogenous hormones from OC on exercise and rehabilitation outcomes is critical for optimizing treatment strategies and promoting equitable care for females. Despite the widespread use of OC (17), their impact on neuromuscular function and recovery remains largely understudied, representing a significant gap in both clinical and exercise science research. The purpose of this study was twofold. First, we aimed to evaluate the effect of OC use on grip strength loss and surface electromyographic (sEMG) excitation—used as an index of neuromuscular activation—following one week of wrist/hand immobilization. Second, we investigated how OC use influenced the weekly recovery trajectory of these outcomes during a home-based rehabilitation program. To broaden the scope of our investigation and place our findings within the broader literature on sex differences, we also included male participants. Given that this study represents a novel line of inquiry, we chose to focus exclusively on monophasic OC users, as this formulation is among the most commonly used (23). While we did not formulate a specific directional hypothesis due to the exploratory nature of the research, we anticipated that females would exhibit greater muscle weakness and slower recovery compared to males, independent of OC use.

## MATERIALS AND METHODS

### Participants

Right-handed adults between the ages of 18 and 35 years were recruited to participate in this study. We recruited three distinct groups: 1) males, 2) females using monophasic OC, and 3) females not using any form of contraceptive (i.e., non-OC). Screening was conducted via telephone prior to scheduling a study visit to confirm eligibility based on the inclusion and exclusion criteria. For inclusion in the OC group, females were required to have used monophasic OC consistently for at least the past six months. Non-OC females were required to self-report regular menstrual cycles, defined as having a period every 21 to 35 days and/or at least 5 periods in the last 6 months. Individuals were excluded if they reported upper extremity pain or discomfort, defined as a score > 2/5 on question 9 or > 1/5 on questions 10 or 11 of the QuickDASH outcome measure (24). Additional exclusion criteria included left-hand dominance (the left wrist/hand was immobilized), body mass index (BMI) outside the range of 18.5–29.9 kg/m^2^, and any history of musculoskeletal injury or surgery involving the elbow, wrist, or hand. Participants were also excluded if they had a diagnosis of a neuromuscular disease (e.g., multiple sclerosis, amyotrophic lateral sclerosis, Parkinson’s disease), metabolic disease (e.g., diabetes, thyroid disorder, metabolic syndrome), or if they had a personal or family history of blood clots. The PHQ4 was used to screen for anxiety and depression, with scores ≥ 3 used as cut points (25). Other exclusion criteria included difficulty controlling muscles, history of cancer, stroke, heart attack, or arthritis, prior or current pregnancy, any type of bodily implant, or the use of hormonal contraceptives other than OC (e.g., intrauterine device, implant) within the past two years.

A total of 41 participants enrolled in this study (12 males, 11 OC females, 18 non-OC females). Of these, 11 participants withdrew. The reported reasons for withdrawal included scheduling conflicts (n=2), difficulty completing daily activities due to immobilization (n=3), menstrual cycle irregularities (n=1), and loss of interest (n=5). A total of 30 healthy young adults completed the study and were included in the final dataset, including 10 males, 10 females using monophasic OC, and 10 females not using hormonal contraceptives (non-OC). The mean ± SD age and BMI of the overall sample were 22.6 ± 3.0 years and 23.1 ± 3.3 kg/m^2^, respectively. Group-specific demographics were as follows: males (age: 24.7 ± 3.6 years; BMI: 24.2 ± 3.6 kg/m^2^), non-OC females (age: 22.1 ± 2.7 years; BMI: 23.3 ± 3.2 kg/m^2^), and OC females (age: 21.0 ± 1.3 years; BMI: 21.8 ± 2.8 kg/m^2^). Physical activity was assessed using the International Physical Activity Questionnaire Short Form (*26*). Most participants were classified as highly active, including 7 males, 9 non-OC females, and 5 OC females. Three males and 1 non-OC female reported moderate activity levels, while 5 OC females were classified as having moderate activity.

### Study Design

This study utilized a repeated measures design to evaluate changes in maximal grip force, RFD, and neuromuscular activation across: pre-immobilization (baseline), post-immobilization, and post-rehabilitation. As shown in **Figure 1**, all participants completed *a minimum of* three laboratory visits at the University of Central Florida (UCF), each spaced exactly seven days apart. The visits included: (1) baseline testing and immobilization initiation, (2) post-immobilization testing following one week of wrist/hand bracing, and (3) post-rehabilitation testing after one week of at-home recovery exercises. Rehabilitation involved twice-daily sessions and continued until each participant’s peak grip force returned to within 2.5% of their baseline value. In other words, participants returned for testing for a fourth or fifth visit if they did not demonstrate recovery. All testing sessions were conducted at the same time of day (±1 hour) to minimize diurnal variation. To promote adherence and consistency, each participant was assigned a designated study investigator who provided frequent support throughout the study. Non-OC females were immobilized during their luteal phase, while OC females did so during the active hormone phase of their pill cycle, avoiding the placebo pill phase (*full details provided below*).

**Figure 1.**
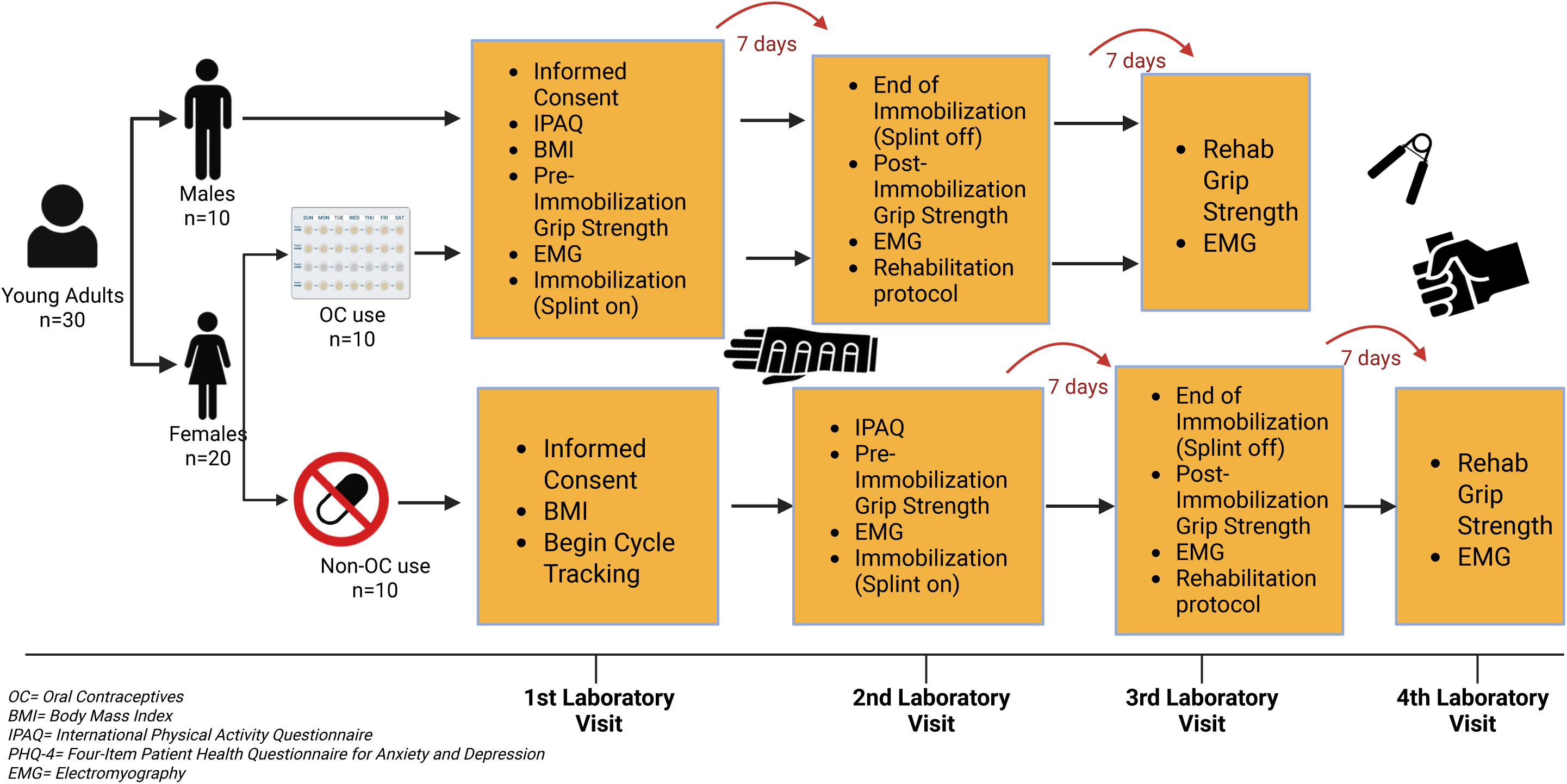
An overview of the experimental design of this study.

Participants were permitted to engage in general physical activity during the immobilization week but were instructed to avoid any upper-body resistance training. All testing procedures were performed by the same trained research personnel to ensure protocol fidelity and reliability. The study was approved by the UCF Institutional Review Board (IRB #6302) and was prospectively registered on ClinicalTrials.gov (ID: NCT06275048). All participants provided written informed consent after reviewing and acknowledging the study procedures and associated risks. Participants were provided a $100 Amazon gift card for completing all study procedures.

### Immobilization

Immediately following the baseline testing visit, participants were fitted with a commercially available wrist and hand splint (Functional Resting Hand Splint, OSK), and began a one-week immobilization period of the left (non-dominant) wrist/hand. The splint featured multiple adjustable straps and was designed to completely restrict movement at the wrist and thumb joints. Although limited motion of the second through fifth digits (index to little finger) was possible, participants were discouraged from finger movement throughout the immobilization period. During splint fitting, additional Velcro straps were provided over the distal interphalangeal joint of digits two through five. A thin stocking was placed between the skin and splint to minimize irritation, and participants were familiarized with proper splint use during the initial visit. They were instructed to wear the splint 24 hours a day for the full 7-day period, including during sleep and personal hygiene routines. To maintain dryness while showering or bathing, participants were provided with a plastic covering and rubber band, along with clear instructions on proper application.

To promote skin hygiene, participants were allowed to replace the underlying stocking midway through the immobilization period but were required to initiate a video call (e.g., FaceTime) with a research team member during the process. This allowed for remote monitoring to ensure the wrist/hand remained immobilized during brief splint removal. Any other instance of brace removal required prior approval from the assigned research team member. Participants were expected to notify the team via video call if removal was necessary, and all instances were documented by both the participant and research staff. Frequent check-ins were conducted by each participant’s assigned compliance officer to ensure comfort, answer questions, and promote adherence.

### Post-Immobilization Rehabilitation

Following the one-week immobilization period, participants began a structured, at-home rehabilitation program designed by physical therapists to restore wrist and hand strength and mobility. Rehabilitation commenced the day after the post-immobilization testing session and continued until the participant’s peak grip force returned to within 2.5% of their baseline value. The program was performed once in the morning and once in the evening, with each session lasting approximately 15 minutes. During the post-immobilization testing session, participants were individually instructed by a member of the research team on proper technique, progression, and expectations for each exercise. A printout was provided to each participant that they could refer to for guidance. Participants remained in regular contact with their assigned research team member, who provided check-ins to ensure compliance, address concerns, and monitor progress.

Each at-home rehabilitation session began with a brief warm-up consisting of wrist circles performed in both clockwise and counterclockwise directions for 30 seconds each. This was followed by a series of strengthening exercises. Participants completed five sets of eight gripper squeezes at approximately 80% of their perceived maximum effort (rated 8 out of 10), holding each squeeze for five seconds and resting one minute between sets. Wrist curls were performed with the arm extended and the palm facing upward; participants slowly curled the wrist through its full range of motion, pausing at the top and bottom for a brief hold (2 sets of 10 repetitions). Wrist extensions were completed in a similar position with the palm of the hand facing downward, again emphasizing slow, controlled movement and a squeeze at each end range (2 sets of 10 repetitions). Finger spreads were performed by moving from a closed fist to a fully spread hand and returning to a fist, repeated for 15 repetitions. The overall goal of the rehabilitation program was to rebuild wrist and hand strength gradually, while ensuring safety, consistency, and adherence throughout the recovery process.

### Menstrual Cycle Monitoring and Hormonal Phase Standardization

To account for potential hormonal influences on neuromuscular function, menstrual cycle tracking and hormonal phase standardization procedures were implemented based on each female participant’s OC use status. Participants were categorized as either using monophasic OC or naturally cycling (non-OC).

#### OC Participants

Participants using monophasic OC were required to be on a 28-day pill regimen, consisting of 21 days of active hormone pills followed by 7 days of placebo pills, for at least six months prior to study enrollment. Common monophasic OC names/brands included: Sprintec, Blisovi, Tyblume, and Loestrin Fe. To standardize hormone exposure during the immobilization period, splinting was initiated during the active hormone phase, specifically between days 1 and 15 of the 21-day active pill cycle. Immobilization was not started during the hormone withdrawal period. Research personnel coordinated closely with each OC participant to verify their pill schedule and ensure that the timing of immobilization aligned with active hormone administration.

#### Non-OC Participants

Female participants not using OC were instructed to monitor their menstrual cycle for at least one month prior to data collection using the *Fertility Friend* mobile app (Ottawa, Canada).

Upon enrollment, participants were provided with a digital thermometer and instructed to record their basal body temperature within the app each morning within one hour upon waking. The *Fertility Friend* app uses temperature data, along with optional entries for symptoms such as cervical fluid consistency, menstruation or spotting, and other physical signs, to estimate the timing of ovulation and identify different phases of the menstrual cycle, including the luteal phase (27). Members of the research team provided one-on-one instruction on how to use the app, including how to enter daily data, interpret cycle tracking results, and ensure data completeness. This allowed the research team to confirm the onset of each participant’s luteal phase and to initiate immobilization during this hormonally consistent window.

### Maximal Grip Force

Grip force was assessed using a hand-held Grip Force Transducer (Model MLT004/ST, ADInstruments, Dunedin, New Zealand). The transducer was connected to a PowerLab data acquisition system and LabChart software (v8.0, ADInstruments) for real-time data collection and analysis, sampled at 2,000 Hz. During testing, participants stood upright and were instructed to maintain a 90-degree elbow angle on the testing side, verified using a goniometer by a member of the research team. Each participant completed five maximal voluntary grip contractions per arm. For each trial, participants were instructed to squeeze the grip transducer as quickly and forcefully as possible and maintain the contraction for three seconds. Strong verbal encouragement to squeeze “hard and fast” was provided during each effort, and participants received real-time visual feedback on a monitor displaying the force trace. A one-minute rest was provided between trials to minimize fatigue. Testing was conducted bilaterally, with the order of arm testing determined by coin flip (heads = right arm first, tails = left arm first). Post-immobilization testing followed the same procedures, with the order of arm testing reversed from baseline. Post-rehabilitation testing followed the same standardized protocol to ensure consistency across sessions.

To complete the study, participants were required to return to within 2.5% of their baseline peak grip force (N) on the immobilized limb. If this threshold was not met, participants continued with at-home rehabilitation for an additional week, followed by a repeat testing session using the same procedures.

### sEMG Excitation

sEMG activity was detected from the extensor carpi radialis brevis (ECRB) and flexor digitorum superficialis (FDS) during all grip force testing using a bipolar electrode configuration with an interelectrode distance of 20 mm. Two surface electrodes were placed over the muscle belly of each target muscle, and a ground electrode was positioned over the C7 spinous process. Data were collected using a PowerLab data acquisition system and LabChart software (v8.0, ADInstruments, Dunedin, New Zealand), sampled at 2,000 Hz.

Electrode placement for the ECRB followed previously published protocols (28). The lateral epicondyle and radial styloid process were used as anatomical landmarks, and electrodes were positioned along the proximal one-third of the lateral forearm. For the FDS, electrode placement was individualized to account for differences in forearm size (29). The distance between the medial epicondyle and ulnar styloid was measured, and electrodes were placed in the distal one-third of the forearm to minimize crosstalk from adjacent wrist muscles. The skin was lightly marked with a permanent marker to guide consistent placement prior to electrode placement. Any hair was shaved to reduce skin impedance, then the skin was cleaned with alcohol wipes. Electrodes were applied bilaterally before the start of testing to ensure efficient transitions between arms during grip assessments.

### Signal Processing

All force and sEMG signals were processed offline using custom-written software developed in LabVIEW (2017, National Instruments, Austin, TX). Force signals were low-pass filtered using a second-order Butterworth filter with a 50 Hz cutoff frequency to reduce high-frequency noise while preserving the transient features critical for RFD analysis. Peak grip force was defined as the highest force value (N) recorded during each 3-second maximal voluntary contraction. RFD was calculated as the linear slope of the filtered force-time curve from contraction onset to 30 and 200 ms and expressed in newtons per second (N/s). These time-specific RFD intervals may reflect distinct neuromuscular characteristics, with early-phase RFD (e.g., 0–30 ms) thought to be more influenced by neural drive and motor unit recruitment, and later-phase RFD (e.g., 0–200 ms) reflecting a greater contribution of muscle contractile properties and maximal strength capacity (30). Contraction onset was manually identified by a single investigator (H.M.B.) through visual inspection of the raw force signal and defined as the point where force began a consistent rise above baseline. Trials with unstable or noisy baselines were excluded from analysis.

sEMG signals were bandpass filtered between 10–500 Hz and analyzed without rectification. The mean value of each signal was removed to eliminate DC offset. Muscle excitation was quantified as the root mean square (RMS) value, in microvolts (μV), across the full 3-second grip contraction.

### Statistical Analyses

For all dependent variables, the values from the three trials with the best performance were averaged and used in the final analysis. Prior to any group comparisons, we objectively determined that that there were no statistical outliers using the absolute deviation around the mean approach described by Leys et al. (31), with a threshold of ±3. Initial analyses showed no meaningful changes in peak grip force for the right (non-immobilized) limb, supporting its role as a control and reducing concerns about cross-education effects. Therefore, to simplify interpretation, all detailed analyses focus on the left (immobilized) limb. All statistical analyses were conducted using linear mixed-effects models to evaluate changes across three time points (Pre, Post, Post-Rehabilitation) and between three groups (males, females not using OC, and females using OC). Each dependent variable (peak force, RFD30, RFD200, sEMG excitation for FDS and ECBR) was modeled with fixed effects for time, group, and their interaction, and a random intercept for participant ID to account for repeated measures. Unexpectedly, three female participants (1 OC, 2 non-OC) showed very small *increases* in peak force following immobilization (mean increase = 1.56%) and were therefore not required to participate in the rehabilitation phase, resulting in missing data for these individuals. Missing data were handled under the assumption of missing at random, and no imputation or last observation carried forward was applied. Model-estimated marginal means and contrasts were used to examine specific differences between posttest and post-rehabilitation values. Effect sizes were estimated as partial eta squared (η^2^_p_), interpreted using conventional thresholds (small ≥ 0.01, moderate ≥ 0.06, large ≥ 0.14) (32). For transparency and interpretability, observed means ± standard deviations (SD) are also reported for each time point. Statistical significance was defined *a priori* as p < 0.05 for all comparisons. All statistical analyses were conducted with JASP software (version 0.19.3, The JASP Team. 2025).

## RESULTS

Table 1 shows the mean±SD values for each of the dependent variables described below for the immobilized wrist/hand. These data are shown for the non-immobilized control wrist/hand in Table 2. The results described below correspond to the immobilized wrist/hand only.

**Table 1.** Mean±SD force and sEMG excitation for each group across the three time points. The results correspond to the immobilized wrist/hand. The post-rehabilitation columns do not include data for the 3 females that did not require rehabilitation (i.e., N=27 rather than N=30).

| Dependent Variable | Pretest |  |  | Posttest |  |  | Post-rehabilitation |  |  |
| --- | --- | --- | --- | --- | --- | --- | --- | --- | --- |
|  | Males | OC | Non-OC | Males | OC | Non-OC | Males | OC | Non-OC |
| Peak Force (N) | 410.1±87.1 | 235.1±45.35 | 257.0±41.0 | 338.9±82.8 | 190.7±81.5 | 181.1±50.1 | 430.2±95.4 | 270.8±40.7 | 242.4±54.5 |
| RFD30 (N/s) | 155.4±112.2 | 188.8±142.3 | 134.1±91.5 | 119.8±74.8 | 107.4±57.5 | 89.7±52.4 | 126.0±67.1 | 145.9±105.0 | 187.0±163.7 |
| RFD200 (N/s) | 819.9±225.3 | 577.7±204.3 | 547.7±121.9 | 682.7±216.0 | 417.3±127.8 | 386.5±226.1 | 810.3±317.8 | 527.3±195.1 | 625.8±210.9 |
| FDS Excitation (μV RMS) | 558.9±277.2 | 460.4±142.0 | 477.4±278.6 | 481.6±247.3 | 368.7±178.1 | 401.1±295.2 | 559.2±253.1 | 656.9±298.9 | 670.1±313.2 |
| ECRB Excitation (μV RMS) | 608.4±351.1 | 405.7±250.6 | 568.6±380.7 | 494.7±332.6 | 430.1±310.3 | 365.2±236.2 | 759.8±501.9 | 523.3±253.8 | 673.5±425.1 |

**Table 2.** Mean±SD force and sEMG excitation for each group across the three time points. The results correspond to the non-immobilized wrist/hand. The post-rehabilitation columns do not include data for the 3 females that did not require rehabilitation (i.e., N=27 rather than N=30).

| Dependent Variable | Pretest |  |  | Posttest |  |  | Post-rehabilitation |  |  |
| --- | --- | --- | --- | --- | --- | --- | --- | --- | --- |
|  | Males | OC | Non-OC | Males | OC | Non-OC | Males | OC | Non-OC |
| Peak Force (N) | 432.4±97.1 | 239.2±55.1 | 287.6±58.4 | 432.9±96.6 | 227.4±59.4 | 273.1±60.1 | 446.7±102.9 | 235.9±58.9 | 272.9±37.6 |
| RFD30 (N/s) | 252.4±104.5 | 131.2±80.4 | 123.4±62.5 | 228.8±171.8 | 142.3±85.7 | 140.9±69.2 | 215.4±122.4 | 136.7±61.8 | 127.5±68.9 |
| RFD200 (N/s) | 1045.7±327.0 | 552.6±239.2 | 614.4±139.4 | 1020.0±269.7 | 601.9±173.4 | 584.8±106.7 | 1021.7±301.4 | 580.8±163.2 | 571.3±189.6 |
| ECRB Excitation (μV RMS) | 611.4±402.2 | 483.3±250.1 | 663.1±551.8 | 621.9±442.0 | 443.6±247.4 | 673.0±412.3 | 584.5±382.2 | 557.8±302.7 | 618.2±368.5 |
| FDS Excitation (μV RMS) | 568.5±321.6 | 474.5±214.3 | 587.8±451.2 | 545.8±217.6 | 428.6±181.7 | 541.7±289.8 | 514.2±211.4 | 539.3±212.6 | 628.1±331.7 |

### Peak Force

Peak force results have been highlighted in **Figure 2** and **Figure 3**. Figure 3 displays the wide variability in peak force responses to immobilization. The results from the linear mixed-effects model indicated no significant time × group interaction (p > 0.20), indicating that changes in peak force over time did not differ significantly between groups. The estimated effect size for the interaction was small/medium (η^2^_p_ = 0.060). Specifically, the difference in the posttest decline between males and females not using OC was 18.98 N (95% CI: –15.96 to 53.92, p = 0.287), and between males and females using OC was 22.78 N (95% CI: –12.16 to 57.72, p = 0.201).

**Figure 2.**
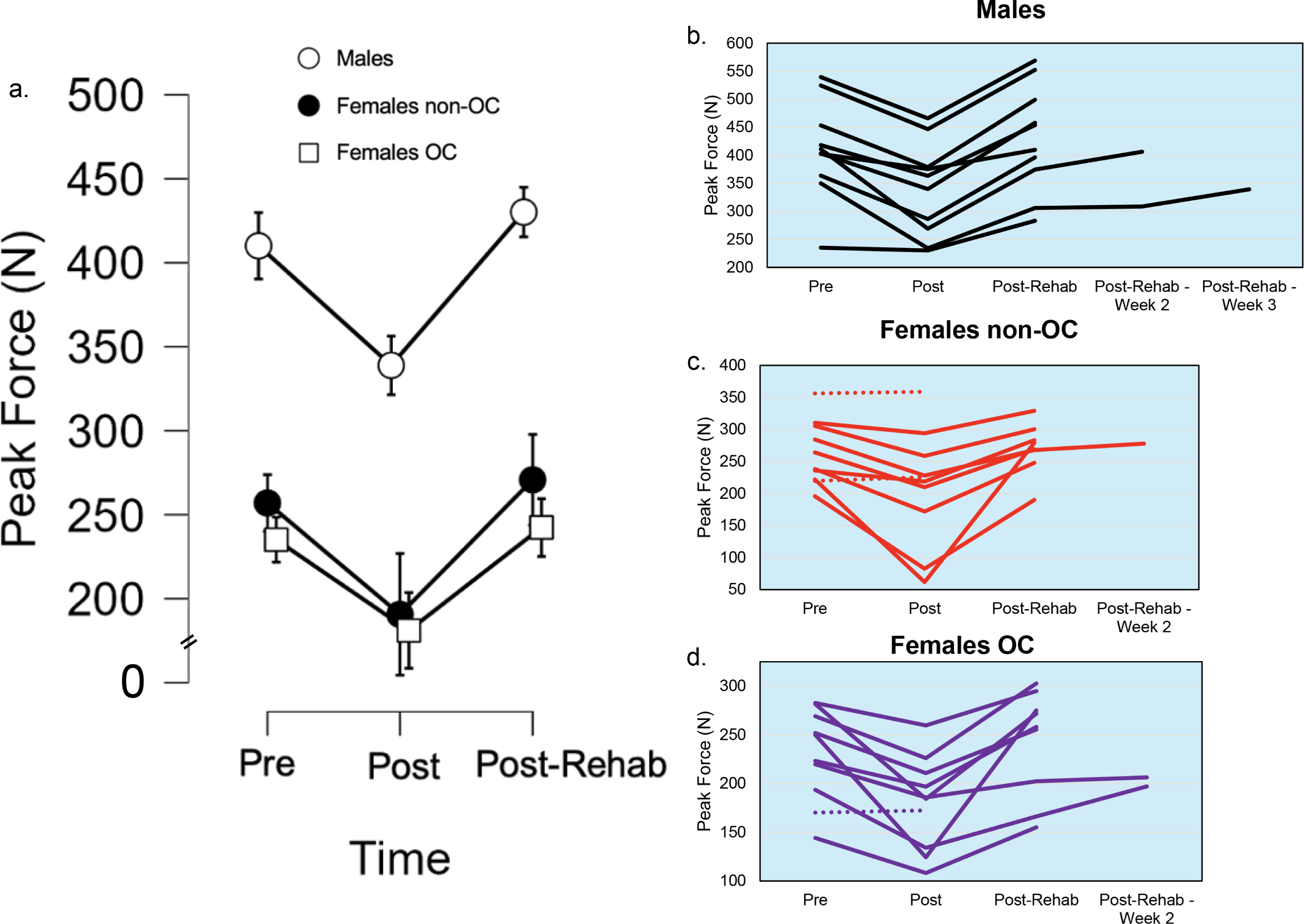
Changes in peak grip force of the immobilized hand. **a.** shows mean ± standard error of the mean peak force at pretest, posttest, and post-rehabilitation. Panels b–d show individual participant responses for each of the three groups. Note the differing number of data points, with dashed lines for those that did not lose grip force, depicting the varying need for rehabilitation based on recovery of peak force back to baseline. Peak force significantly declined posttest and returned to or exceeded baseline following rehabilitation (p < 0.001). Dashed lines correspond to participants that did not require rehabilitation. The five lines that extend beyond the Post-Rehab time point represent participants who required extended rehabilitation and returned for additional follow-up testing.

**Figure 3.**
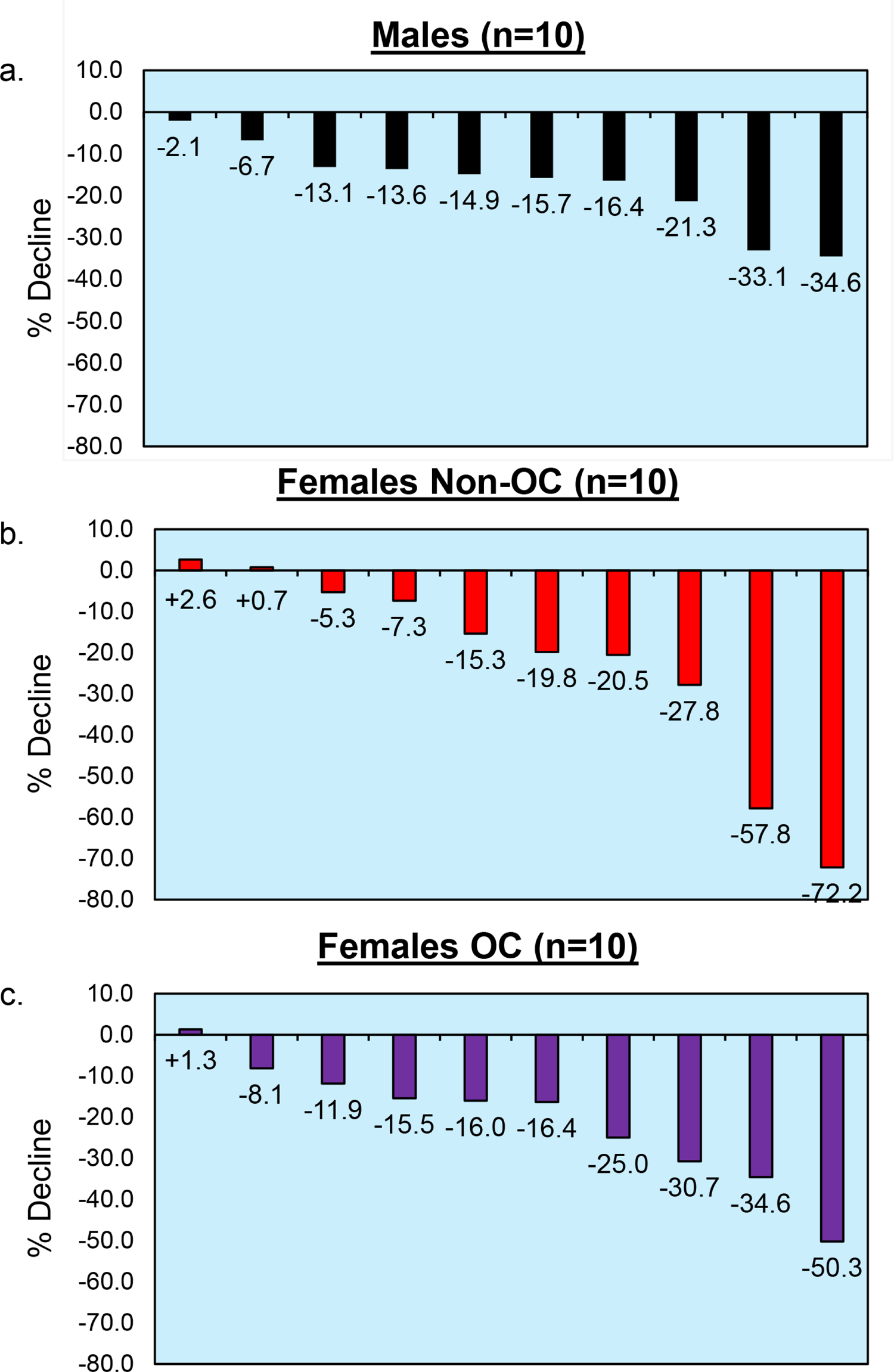
Percent decline in peak grip force following immobilization by group. (a–c) Individual percent changes in peak grip force from pretest to posttest are shown for males (a), females not using OC (b), and females using OC (c). Group means ± SD are: males = –17.2 ± 10.3%, non-OC = –22.3 ± 24.7%, OC = –20.7 ± 14.8%. There were no statistically significant group differences in the magnitude of decline (p = 0.545), indicating a similar response to immobilization across sex and contraceptive status.

Main effects of group revealed that, when collapsing across time, females not using OC demonstrated 146.93 N (95% CI: –208.54 to –85.32, p < 0.001) lower peak force than males, while females using OC exhibited 181.48 N (95% CI: –243.09 to –119.87, p < 0.001) lower peak force. The effect of group was large (η^2^_p_ = 0.421), indicating robust sex differences in peak force.

A significant main effect of time was also observed (η^2^_p_ = 0.362). Peak force declined from pretest to posttest by 71.19 N (95% CI: –95.90 to –46.49, p < 0.001) across all groups. Although the average peak force at post-rehabilitation was 20.04 N higher than at pretest, this difference was not statistically significant (95% CI: –4.66 to 44.75, p = 0.112). However, a direct contrast between posttest and post-rehabilitation values revealed a significant increase in force (91.24 ± 12.61 N, t(78) = 7.24, p < 0.0001), indicating robust recovery following the rehabilitation period across all groups.

### RFD30

RFD30 results have been highlighted in **Figure 4**. The results from the linear mixed-effects model indicated no significant time × group interactions (p > 0.25 for all contrasts), indicating that changes in RFD30 over time did not differ significantly between groups. The estimated effect size for the interaction was small (η^2^_p_ = 0.041). The difference in post-rehabilitation recovery between males and females not using OC was +56.75 N/s (95% CI: –40.72 to 154.22, p = 0.254), while the difference between males and females using OC was –7.16 N/s (95% CI: –96.84 to 82.51, p = 0.876).

**Figure 4.**
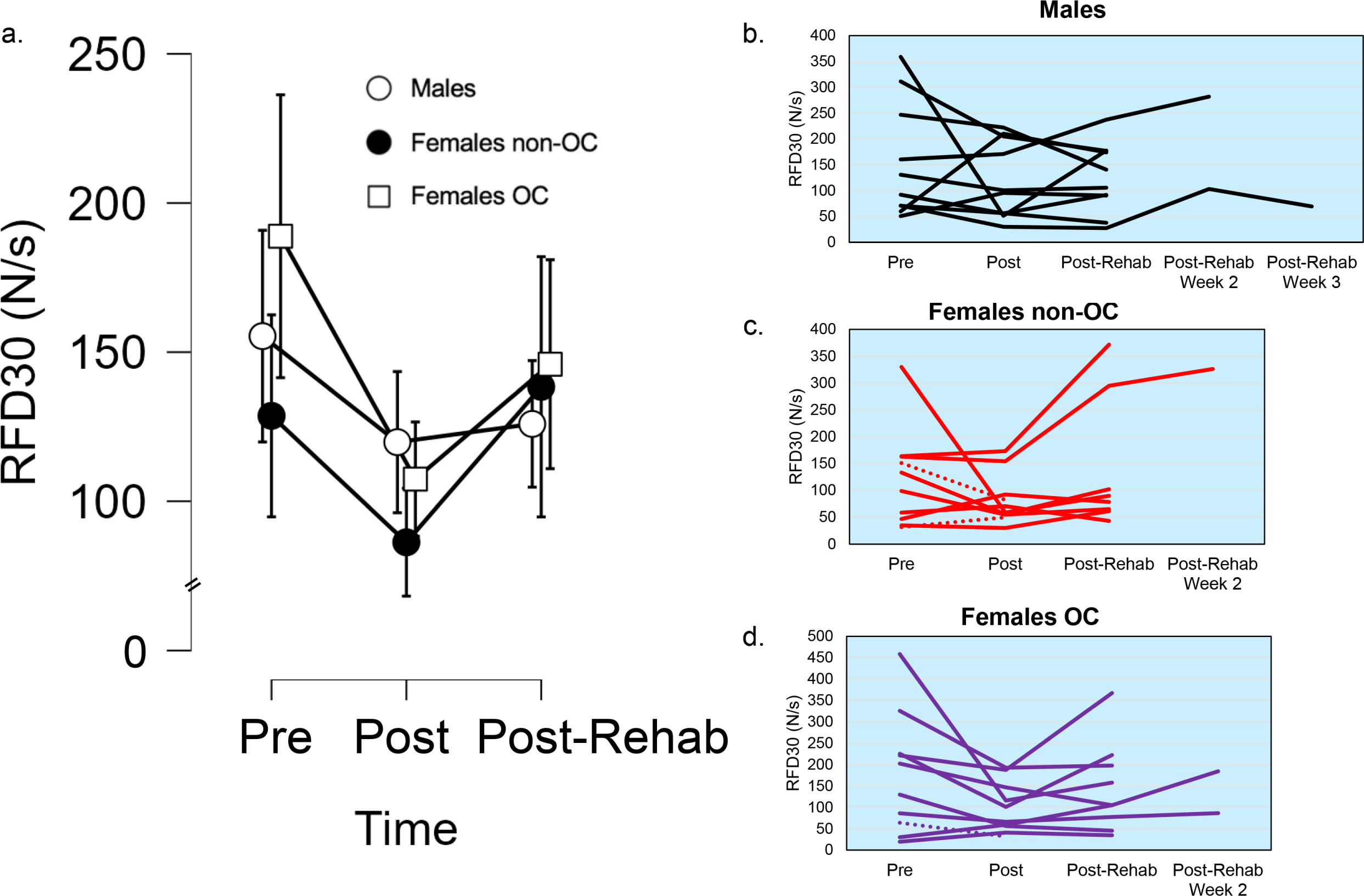
RFD measured at 30 ms. (a) Group mean ± standard error RFD30 values across time points. (b– d) Individual trajectories for males (b), non-OC females (c), and OC females (d). No significant time effect was observed (p = 0.263), and recovery post-rehabilitation was incomplete. RFD30 exhibited greater inter-individual variability and lower responsiveness compared to peak force and RFD200. Dashed lines correspond to participants that did not require rehabilitation. The five lines that extend beyond the Post-Rehab time point represent participants who required extended rehabilitation and returned for additional follow-up testing.

When collapsing across time points, no significant main effects of group were detected. Compared to males, females not using OC had 34.17 N/s lower RFD30 (95% CI: –117.74 to 49.40, p = 0.423), while females using OC showed a non-significant increase of 20.99 N/s (95% CI: –62.58 to 104.56, p = 0.622). The overall effect of group was small (η^2^_p_ = 0.058).

Time effects revealed that RFD30 decreased from pretest to posttest (150.98 ± 114.18 N/s vs. 100.94 ± 62.22 N/s), with a model-estimated mean reduction of 35.54 N/s (95% CI: –97.81 to 26.73, p = 0.263). Following rehabilitation, average RFD30 increased to 141.46 ± 97.49 N/s, reflecting a return toward pretest values. However, the model-estimated difference between posttest and post-rehabilitation values was not statistically significant (difference = +6.12 ± 31.77 N/s, t(75) = 0.19, p = 0.848), and the time effect overall was small (η^2^_p_ = 0.051). These findings suggest that although mean RFD30 improved after rehabilitation, variability in individual responses may have limited the ability to detect a statistically significant change.

### RFD200

RFD30 results have been highlighted in **Figure 5**. The results from the linear mixed-effects model indicated no significant time × group interactions (all p > 0.44), suggesting that changes in RFD200 over time did not differ by group. The estimated effect size for the interaction was small (η^2^_p_ = 0.043). For example, the group difference in recovery between posttest and post-rehab was +66.26 N/s for females not using OC (95% CI: –104.03 to 236.56, p = 0.446) and –32.31 N/s for females using OC (95% CI: – 188.20 to 123.58, p = 0.685).

**Figure 5.**
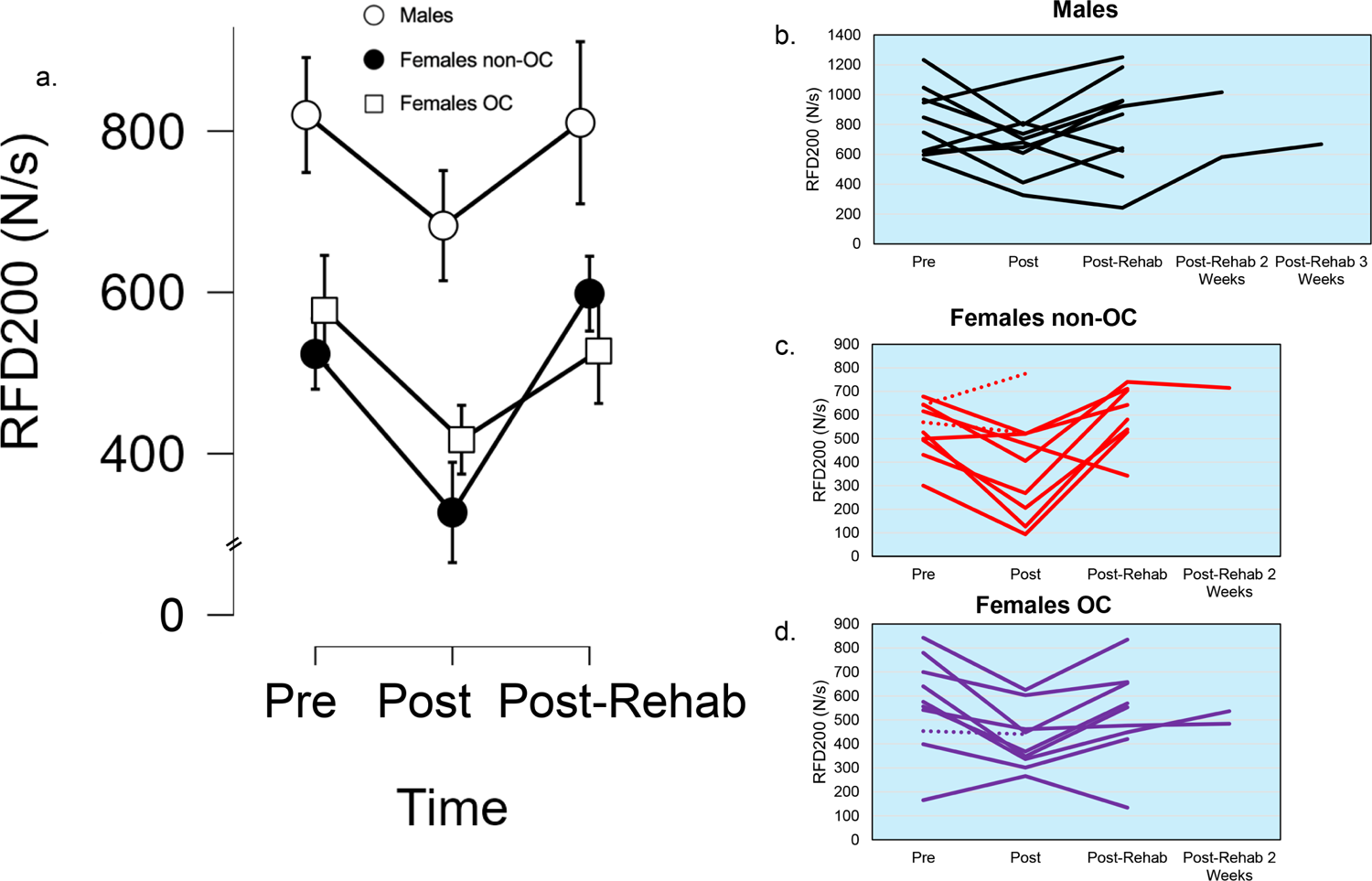
RFD measured at 200 ms. (a) Mean ± standard error values for RFD200 at pretest, posttest, and post-rehabilitation. (b–d) Individual data plotted for males (b), non-OC females (c), and OC females (d). A significant time effect was found (p = 0.013), with posttest RFD200 reduced and recovery observed following rehabilitation (post-rehab > posttest, p = 0.024). Dashed lines correspond to participants that did not require rehabilitation. The five lines that extend beyond the Post-Rehab time point represent participants who required extended rehabilitation and returned for additional follow-up testing.

Main effects of group revealed that, when collapsing across time, females not using OC demonstrated 279.77 N/s lower RFD200 than males (95% CI: –458.50 to –101.04, p = 0.002), and females using OC demonstrated 254.58 N/s lower RFD200 than males (95% CI: –433.31 to –75.85, p = 0.005). The group effect was moderate-to-large (η^2^_p_ = 0.084), indicating substantial sex differences (males>females) in RFD200 regardless of time point.

Time effects indicated a significant reduction in RFD200 from pretest (641.73 ± 220.05 N/s) to posttest (505.91 ± 225.70 N/s), with a model-estimated drop of 137.17 N/s (95% CI: –245.34 to –29.00, p = 0.013). Following rehabilitation, mean RFD200 increased to 648.80 ± 269.33 N/s, nearly identical to pretest levels. A direct comparison between posttest and post-rehabilitation values revealed a statistically significant increase of 127.59 ± 55.19 N/s (t(75) = 2.31, p = 0.024), supporting the effectiveness of the rehabilitation period in restoring RFD200. The effect of time was moderate (η^2^_p_ = 0.073).

### FDS Excitation

FDS excitation results have been highlighted in **Figure 6**. The results from the linear mixed-effects model indicated no significant time × group interactions (all p > 0.05), though the post-rehabilitation increase in females appeared greater than in males. The estimated interaction effect for females not using OC was +203.93 µV (95% CI: –15.61 to 423.48, p = 0.069), while the effect for females using OC was +210.11 µV (95% CI: –4.50 to 424.71, p = 0.055), representing small-to-moderate interaction effects (η^2^_p_ = 0.061).

**Figure 6.**
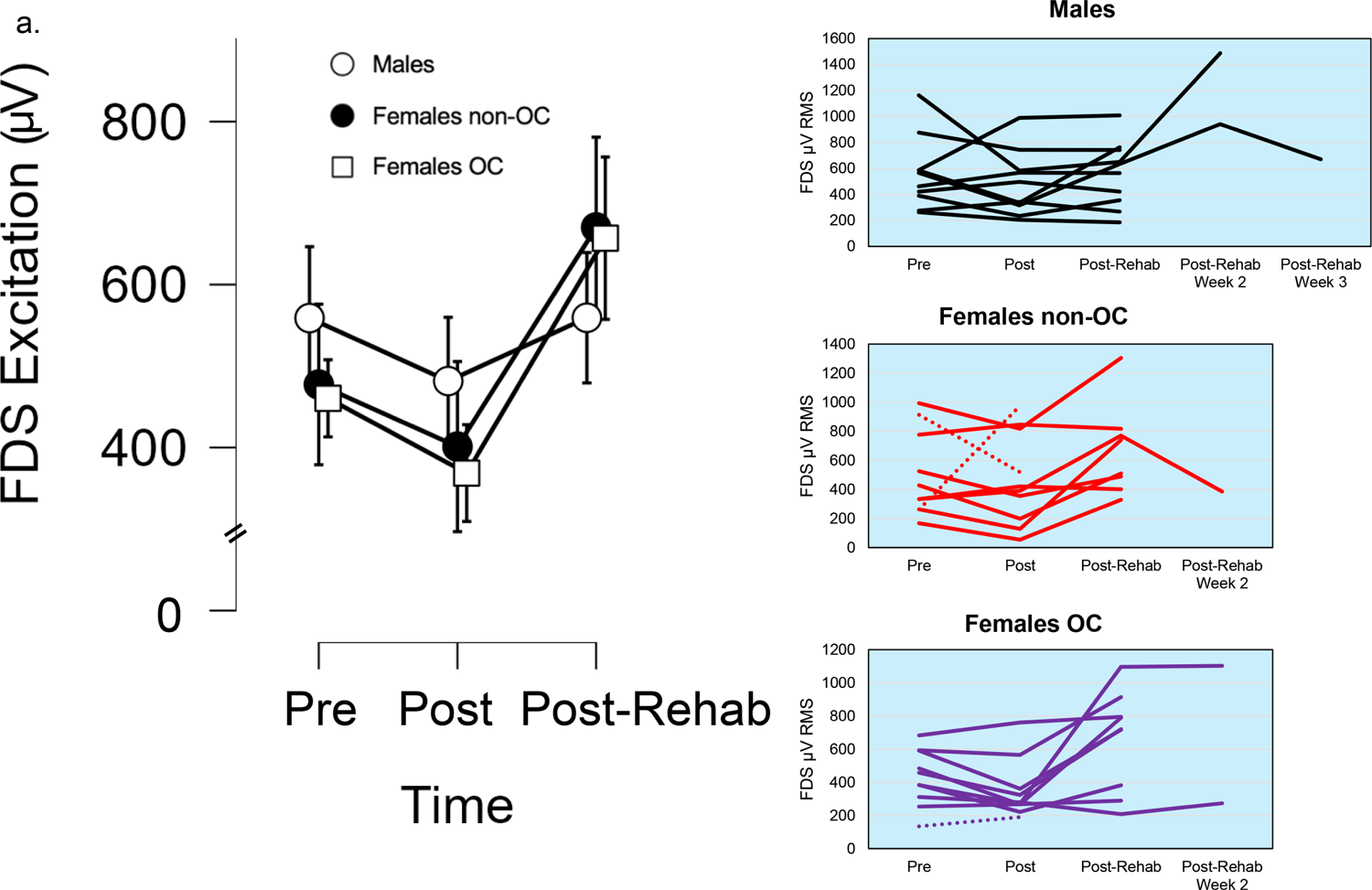
sEMG excitation of the FDS muscle. (a) Mean ± standard error values for sEMG excitation (μV RMS) across time points. (b–d) Individual participant data for males (b), non-OC females (c), and OC females (d). FDS excitation increased following rehabilitation, with values returning near or above baseline. Although the change was not statistically significant (p = 0.062), substantial inter-individual variability was observed, suggesting diverse neuromuscular responses to disuse and recovery. Dashed lines correspond to participants that did not require rehabilitation. The five lines that extend beyond the Post-Rehab time point represent participants who required extended rehabilitation and returned for additional follow-up testing.

Main effects of group were not significant. Compared to males, females not using OC exhibited 59.40 µV lower excitation (95% CI: –290.66 to 171.86, p = 0.615), while females using OC exhibited 131.07 µV lower excitation (95% CI: –362.33 to 100.19, p = 0.267). The group effect was small (η^2^_p_ = 0.049).

Time effects revealed that average FDS excitation declined from pretest (495.37 ± 249.78 µV) to posttest (434.26 ± 252.21 µV), though this reduction was not statistically significant (–77.28 µV, 95% CI: –226.20 to 71.64, p = 0.309). Excitation values increased following rehabilitation to 624.62 ± 280.57 µV, exceeding pretest levels. However, the model-estimated increase from posttest to post-rehab (+77.65 ± 75.98 µV, t(78) = 1.02, p = 0.310) was not statistically significant. The overall time effect was small (η^2^_p_ = 0.056), suggesting a trend toward increased muscle activation following rehabilitation that did not reach significance due to variability in individual responses.

### ECBR Excitation

ECBR excitation results have been highlighted in **Figure 7**. The results from the linear mixed-effects model indicated no significant time × group interactions (all p > 0.39), suggesting that changes in ECBR excitation over time did not differ by group. The interaction effect for females not using OC was +11.39 µV (95% CI: –264.45 to 287.24, p = 0.935), and for females using OC was –20.16 µV (95% CI: –289.75 to 249.43, p = 0.883), indicating minimal group-specific differences. The estimated effect size for the interaction was small (η^2^_p_ = 0.059).

**Figure 7.**
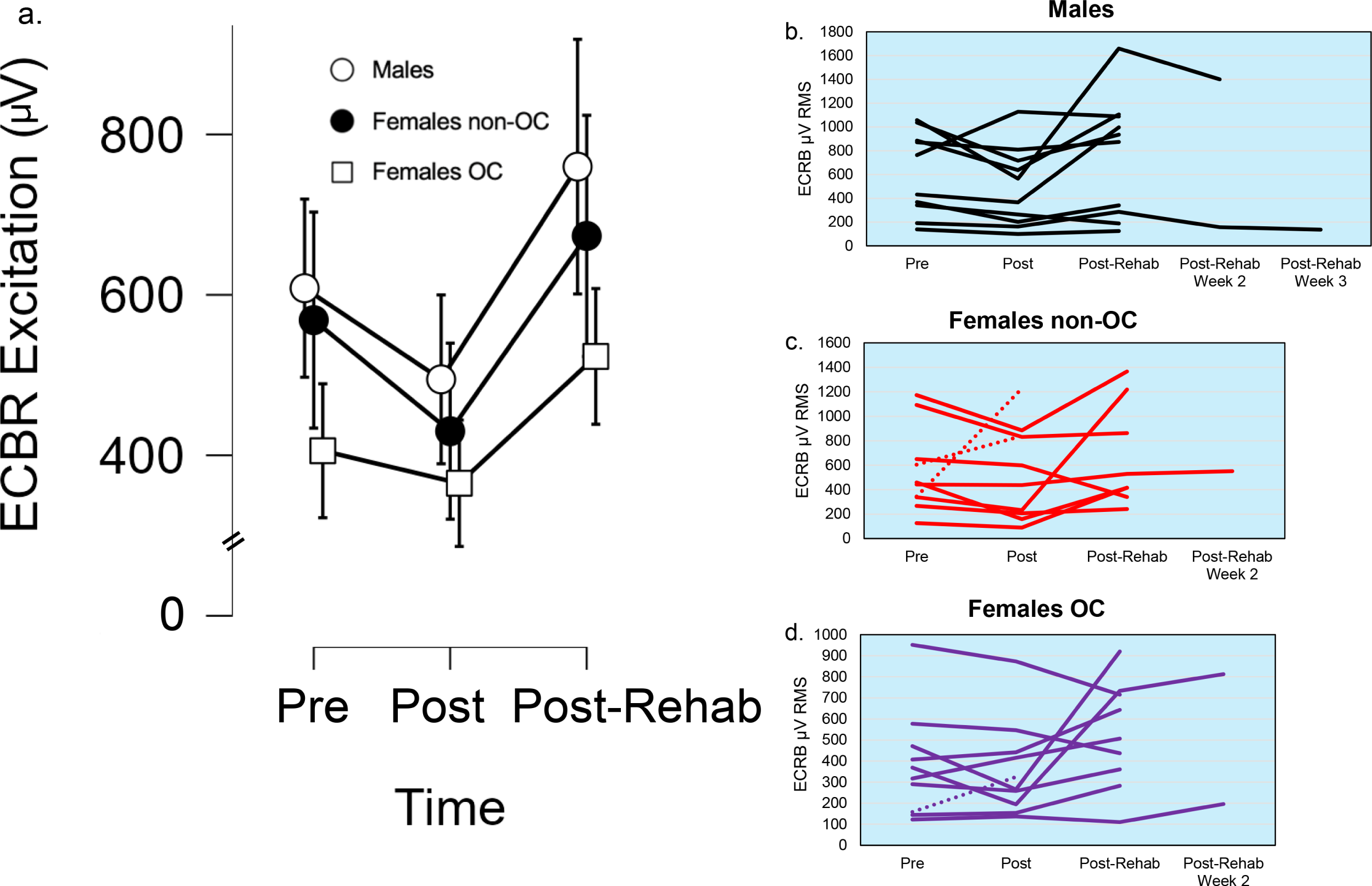
sEMG excitation of the ECRB muscle. (a) Mean ± standard error values for sEMG excitation (μV RMS) across time points. (b–d) Individual participant data for males (b), non-OC females (c), and OC females (d). ECRB excitation significantly increased following rehabilitation (p = 0.007), with post-rehab values exceeding both pretest and posttest levels. This suggests enhanced volitional drive or neuromuscular re-engagement during the recovery phase. Dashed lines correspond to participants that did not require rehabilitation. The five lines that extend beyond the Post-Rehab time point represent participants who required extended rehabilitation and returned for additional follow-up testing.

Main effects of group were also nonsignificant. Females not using OC had 58.62 µV lower ECBR excitation than males (95% CI: –363.70 to 246.46, p = 0.706), and females using OC had 227.42 µV lower excitation (95% CI: –532.50 to 77.66, p = 0.144). The group effect was small (η^2^_p_ = 0.048).

Mean ECBR excitation slightly declined from pretest (513.03 ± 322.08 µV) to posttest (468.83 ± 319.66 µV), with a model-estimated reduction of 113.66 µV (95% CI: –300.72 to 73.40, p = 0.234). Following rehabilitation, excitation increased to 655.41 ± 407.42 µV. The model-estimated difference between posttest and post-rehabilitation was significant (difference = +265.10 ± 95.44 µV, t(78) = 2.78, p = 0.007), indicating a substantial increase in muscle activation following rehabilitation. The effect of time was moderate (η^2^_p_ = 0.081), highlighting meaningful neuromuscular recovery during the post-rehabilitation phase.

## DISCUSSION

Several studies have reported that females lose greater muscle strength (8–12) and/or exhibit slower recovery (12,13) following limb immobilization compared to males. However, the mechanistic contributors to these sex-based differences remain poorly understood. The key findings of the present study were that: (1) use of monophasic OC did not influence the decline in grip force following one week of wrist/hand immobilization, and (2) OC use did not alter recovery trajectories during a subsequent at-home rehabilitation program. Additionally, we observed broadly similar patterns of decline and recovery among males and females, regardless of OC status, though individual responses to immobilization were variable (Figure 3). Below, we outline the significance, implications, strengths, and limitations of this work, as well as provide recommendations for future research.

A novel finding of the present study was that monophasic OC use did not influence peak or rapid force loss, nor did it alter neuromuscular activation of the FDS or ECRB muscles following one week of wrist/hand immobilization. Despite prior hypotheses from our group suggesting that exogenous hormones may blunt or exacerbate neuromuscular declines during disuse (12), our data revealed no meaningful differences between OC users and non-users across any time point. Specifically, the magnitude of decline in peak grip force and rate of force development at both early (30 ms) and later (200 ms) phases was comparable between groups. Likewise, sEMG excitation for both FDS and ECRB showed similar patterns of post-immobilization suppression and post-rehabilitation enhancement, independent of OC status. These findings suggest that, under the conditions of this study, hormonal contraception does not significantly modulate the neuromuscular response to short-term disuse. This is particularly important given the high prevalence of OC use among young females. Our work provides initial reassurance that neuromuscular function and rehabilitation potential are not compromised by exogenous hormonal regulation. While we acknowledge that peak force responses during immobilization were variable, this variability was observed among all groups, suggesting that OC use or sex were not responsible. Future studies are needed to understand why some participants observe such dramatic declines in peak force during disuse, whereas others fair quite well.

A second noteworthy finding of this study was that OC use did not influence recovery trajectories following wrist/hand immobilization. Our study was, in part, motivated by prior work from Clark et al. (13) and Girts et al. (12), which reported that females may recover more slowly than males after short-term disuse. In contrast, our data did not reveal any differences in recovery between OC users and non-users, nor between females and males, suggesting that monophasic OC use does not negatively affect rehabilitation outcomes. Notably, the majority of participants in all groups required only one week of at-home rehabilitation to return to baseline grip force levels, demonstrating a robust and rapid recovery response. However, five participants required additional weeks of rehabilitation before achieving full recovery, underscoring the presence of inter-individual variability. These individuals were distributed across groups and did not cluster by sex or hormonal status, further reinforcing the conclusion that OC use did not systematically influence recovery duration or magnitude. Collectively, these findings support the clinical interpretation that monophasic OC use does not impair neuromuscular recovery following disuse, and they challenge the idea that hormonal contraception necessitates modified rehabilitation protocols for otherwise healthy young females.

It is important to emphasize that our primary outcome measure was grip peak force, which served as our principal marker of disuse-induced impairment and recovery, and was used to guide decisions regarding rehabilitation progression. This measure was selected for its clinical relevance, ease of interpretation, and direct link to functional performance (33). However, our inclusion of RFD30, RFD200, and sEMG excitation from the FDS and ECBR provided a more comprehensive assessment of neuromuscular function. These outcomes offered distinct, partially independent perspectives on how disuse and rehabilitation influence the neuromuscular system. For example, RFD30 captured very early-phase force production, which was highly variable, similar between sexes, and less responsive to rehabilitation. RFD200 showed more consistent patterns of recovery, aligning more closely with peak force. Similarly, sEMG responses differed between muscles: ECBR excitation significantly increased after rehabilitation, while FDS showed a non-significant trend toward improvement. These nuanced differences suggest that changes in force output and neuromuscular activation do not always occur in parallel, and that reliance on a single outcome measure may overlook important aspects of physiological adaptation. As such, our multidimensional approach provides richer insight into the mechanisms underlying recovery and highlights the potential utility of pairing performance-based and electrophysiological markers in future disuse studies.

Three key decisions made during study design have important implications for contextualizing our findings. First, we elected to focus on users of monophasic OC, given their widespread use (23) and steady hormonal profile across the cycle. This decision enhanced interpretability and reduced variability related to fluctuating hormone concentrations, which are more difficult to account for in biphasic or triphasic formulations. However, this focus limits generalizability to all contraceptive users and excludes potentially important comparisons involving different hormonal delivery mechanisms (e.g., IUDs, patches, injectables). Second, we selected the wrist/hand as our disuse model. Most prior immobilization research has examined the quadriceps (1,34). In contrast, the wrist/hand represents a smaller, non–weight-bearing muscle group, making it a valuable model for examining neural control and precise force generation. This choice allowed us to use grip strength as a reliable, clinically meaningful outcome, but it is possible that disuse and recovery dynamics differ between upper and lower extremities due to variations in fiber type distribution, habitual loading, and neural drive. Third, our sample consisted exclusively of healthy, young adults. While this reduced confounding from comorbidities and age-related variability, it also restricts the clinical translatability of our results. In particular, future research is urgently needed in middle-aged and older adults to examine how aging and menopause influence disuse-related neuromuscular changes and rehabilitation potential. Although some previous studies have evaluated older males (35,36), relatively little is known about the time course, magnitude, and mechanisms of neuromuscular impairment and recovery in older females—a population that may experience compounded risk due to both age and hormonal changes (37).

Ultimately, broadening study designs to include a more diverse participant pool and a variety of disuse models will be critical for advancing sex-informed and age-responsive rehabilitation science.

This study has several strengths that advance our understanding of neuromuscular recovery following short-term disuse. For example, we included both male and female participants, allowing for exploration of sex-based differences, and further distinguished hormonal status among females by evaluating OC use. Unlike many prior immobilization studies that focus solely on muscle loss or early disuse effects, we also examined recovery following a structured rehabilitation period—an important yet understudied phase in the disuse paradigm. Additionally, we incorporated neuromuscular assessments beyond maximal force, including RFD and sEMG, which offer new insights into potential mechanisms of recovery. We also used the non-immobilized limb as a within-subject control to strengthen internal validity (38) and confirm that cross-limb effects did not influence our findings (39). Despite these strengths, some limitations warrant discussion. Menstrual cycle phase was estimated based on self-report and use of the *Fertility Friend* app rather than verified through hormonal assays. While our approach did include an objective, daily measure of basal body temperature, which offers advantages (40), it introduced uncertainty regarding the influence of ovarian hormones—an increasingly acknowledged methodological concern in exercise research (41). We also did not assess muscle mass or architecture, limiting our ability to determine whether changes in performance were due to atrophy, neural factors, or both. While our use of a removable brace may not fully replicate the effects of rigid casting, the substantial force loss observed suggests that the intervention was sufficiently potent. The sample size, though modest, is comparable to other immobilization studies (8,9,42) and was sufficient to detect time-based effects. While additional participants would have strengthened the study, none of our time × group interaction effect sizes indicated meaningful differences that would have been detected with slightly more participants. Nonetheless, our findings should be considered proof-of-concept, and future studies are needed to replicate these results using larger cohorts, hormone verification protocols, and outcome assessments that include both muscle mass and function.

In summary, this study adds to the growing body of literature supporting the inclusion of females in disuse and rehabilitation research, with particular attention to hormonal status. Our findings indicate that the use of monophasic OC had little to no influence on strength loss during immobilization or on neuromuscular recovery following rehabilitation. Recovery trajectories were remarkably similar between males and females, regardless of OC use, suggesting that hormonal contraception does not impair the capacity to respond to short-term disuse or benefit from rehabilitation. Although modest individual variability was observed, these results offer encouraging evidence that clinicians and practitioners need not be concerned about OC use negatively affecting neuromuscular outcomes during brief periods of inactivity or recovery. Future studies with larger samples, hormonal verification, and multimodal outcome measures will be valuable for confirming these findings and refining individualized rehabilitation strategies.

## DATA AVAILABILITY

Source data for this study are available by reasonable request to the corresponding author.

## DISCLAIMERS

The content is solely the responsibility of the authors and does not necessarily represent the official views of the University of Central Florida, Kansas State University, or the University of North Carolina - Chapel Hill.

## AUTHOR CONTRIBUTIONS

Conceived and designed research: All authors; Performed experiments: MSS, HMB, MFE, KES, EJP; Analyzed data: MSS, HMB, MFE, KES, RMR; Interpreted results of experiments: MSS, HMB, MFE, KES, RMR; Prepared figures: MSS, HMB, MFE, KES, EJP; Drafted manuscript: MSS, HMB, MFE, KES, RMR; Edited and revised manuscript: All authors; Approved final version of manuscript: All authors

## REFERENCES

1. Campbell M, Varley-Campbell J, Fulford J, Taylor B, Mileva KN, Bowtell JL. Effect of Immobilisation on Neuromuscular Function In Vivo in Humans: A Systematic Review. Sports Med. 2019 Jun;49(6):931–50.

2. Di Girolamo FG, Fiotti N, Milanović Z, Situlin R, Mearelli F, Vinci P, et al. The Aging Muscle in Experimental Bed Rest: A Systematic Review and Meta-Analysis. Front Nutr. 2021 Aug 4;8:633987.

3. Clark BC, Mahato NK, Nakazawa M, Law TD, Thomas JS. The power of the mind: the cortex as a critical determinant of muscle strength/weakness. Journal of Neurophysiology. 2014 Dec 15;112(12):3219–26.

4. MacLennan RJ, Ogilvie D, McDorman J, Vargas E, Grusky AR, Kim Y, et al. The time course of neuromuscular impairment during short-term disuse in young women. Physiological Reports. 2021;9(1):e14677.

5. Seo F, Clouette J, Huang Y, Potvin-Desrochers A, Lajeunesse H, Parent-L’Ecuyer F, et al. Changes in brain functional connectivity and muscle strength independent of elbow flexor atrophy following upper limb immobilization in young females. Experimental Physiology. 2024;109(9):1557–71.

6. Plotkin DL, Mattingly ML, Anglin DA, Michel JM, Godwin JS, McIntosh MC, et al. Skeletal muscle myosin heavy chain fragmentation as a potential marker of protein degradation in response to resistance training and disuse atrophy. Exp Physiol. 2024 Oct;109(10):1739–54.

7. Michel JM, Godwin JS, Plotkin DL, McIntosh MC, Mattingly ML, Agostinelli PJ, et al. Effects of leg immobilization and recovery resistance training on skeletal muscle-molecular markers in previously resistance-trained versus untrained adults. Journal of Applied Physiology. 2025 Feb;138(2):450–67.

8. Yasuda N, Glover EI, Phillips SM, Isfort RJ, Tarnopolsky MA. Sex-based differences in skeletal muscle function and morphology with short-term limb immobilization. J Appl Physiol (1985). 2005 Sep;99(3):1085–92.

9. Deschenes MR, McCoy RW, Holdren AN, Eason MK. Gender influences neuromuscular adaptations to muscle unloading. Eur J Appl Physiol. 2009 Jan 6;105(6):889.

10. Deschenes MR, McCoy RW, Mangis KA. Factors Relating to Gender Specificity of Unloading-induced Declines in Strength. Muscle Nerve. 2012 Aug;46(2):210–7.

11. Coffey VG, McGlory C, Phillips SM, Doering TM. Does initial skeletal muscle size or sex affect the magnitude of muscle loss in response to 14 days immobilization? Appl Physiol Nutr Metab. 2023 May 1;48(5):411–6.

12. Girts RM, Harmon KK, Rodriguez G, Beausejour JP, Pagan JI, Carr JC, et al. Sex differences in muscle-quality recovery following one week of knee joint immobilization and subsequent retraining. Appl Physiol Nutr Metab. 2024 Jun 1;49(6):805–17.

13. Clark BC, Manini TM, Hoffman RL, Russ DW. Restoration of Voluntary Muscle Strength After 3 Weeks of Cast Immobilization is Suppressed in Women Compared With Men. Archives of Physical Medicine and Rehabilitation. 2009 Jan;90(1):178–80.

14. Kale MB, Wankhede NL, Goyanka BK, Gupta R, Bishoyi AK, Nathiya D, et al. Unveiling the Neurotransmitter Symphony: Dynamic Shifts in Neurotransmitter Levels during Menstruation. Reprod Sci. 2025 Jan;32(1):26–40.

15. Ansdell P, Brownstein C, Škarabot J, Hicks K, Simoes D, Thomas K, et al. Menstrual cycle-associated modulations in neuromuscular function and fatigability of the knee extensors in eumenorrheic women. Journal of Applied Physiology. 2019 Mar 7;126.

16. Piasecki J, Guo Y, Jones EJ, Phillips BE, Stashuk DW, Atherton PJ, et al. Menstrual Cycle Associated Alteration of Vastus Lateralis Motor Unit Function. Sports Med Open. 2023 Oct 24;9(1):97.

17. Kavanaugh ML, Jerman J. Contraceptive method use in the United States: trends and characteristics between 2008, 2012 and 2014. Contraception. 2018 Jan 1;97(1):14–21.

18. Daniels K. Current Contraceptive Status Among Women Aged 15–49: United States, 2017–2019. 2020;(388).

19. Nolan D, McNulty KL, Manninen M, Egan B. The Effect of Hormonal Contraceptive Use on Skeletal Muscle Hypertrophy, Power and Strength Adaptations to Resistance Exercise Training: A Systematic Review and Multilevel Meta-analysis. Sports Med. 2024 Jan;54(1):105–25.

20. Riechman SE, Lee CW. Oral Contraceptive Use Impairs Muscle Gains in Young Women. The Journal of Strength & Conditioning Research. 2022 Nov;36(11):3074.

21. Oxfeldt M, Dalgaard LB, Jørgensen EB, Johansen FT, Dalgaard EB, Ørtenblad N, et al. Molecular markers of skeletal muscle hypertrophy following 10 wk of resistance training in oral contraceptive users and nonusers. J Appl Physiol (1985). 2020 Dec 1;129(6):1355–64.

22. Engstad MK, Seynnes O, Vesterhus I, Hesseberg E, Fjeldberg K, Carlsen MH, et al. Effect of Oral Contraceptive Use on Muscle Hypertrophy Following Strength Training. Scand J Med Sci Sports. 2025 Apr;35(4):e70052.

23. Hall KS, Trussell J. Types of combined oral contraceptives used by U.S. women. Contraception. 2012 Dec;86(6):659–65.

24. Beaton DE, Wright JG, Katz JN. Development of the QuickDASH: Comparison of Three Item-Reduction Approaches. The Journal of Bone & Joint Surgery. 2005 May;87(5):1038–46.

25. Kroenke K, Spitzer RL, Williams JBW, Lowe B. An Ultra-Brief Screening Scale for Anxiety and Depression: The PHQ-4. Psychosomatics. 2009 Nov 1;50(6):613–21.

26. Craig CL, Marshall AL, Sjostrom M, Bauman AE, Booth ML, Ainsworth BE, et al. International Physical Activity Questionnaire: 12-Country Reliability and Validity. Medicine & Science in Sports & Exercise. 2003 Aug;35(8):1381–95.

27. Cabre HE, Joniak KE, Ladan AN, Moore SR, Blue MNM, Pietrosimone BG, et al. Effects of Hormonal Contraception and the Menstrual Cycle on Maximal Strength and Power Performance. Medicine & Science in Sports & Exercise. 2024 Dec;56(12):2385–93.

28. Sorbie GG, Williams MJ, Boyle DW, Gray A, Brouner J, Gibson N, et al. Intra-session and Inter-day Reliability of the Myon 320 Electromyography System During Sub-maximal Contractions. Frontiers in Physiology [Internet]. 2018 [cited 2024 Feb 12];9. Available from: https://www.frontiersin.org/journals/physiology/articles/10.3389/fphys.2018.00309

29. Kawala-Janik A, Podpora M, Baranowski J, Bauer W, Pelc M. Innovative approach in analysis of EEG and EMG signals — Comparision of the two novel methods. In: 2014 19th International Conference on Methods and Models in Automation and Robotics (MMAR) [Internet]. 2014 [cited 2025 Apr 23]. p. 804–7. Available from: https://ieeexplore.ieee.org/document/6957459

30. Maffiuletti NA, Aagaard P, Blazevich AJ, Folland J, Tillin N, Duchateau J. Rate of force development: physiological and methodological considerations. Eur J Appl Physiol. 2016 Jun;116(6):1091–116.

31. Leys C, Ley C, Klein O, Bernard P, Licata L. Detecting outliers: Do not use standard deviation around the mean, use absolute deviation around the median. Journal of Experimental Social Psychology. 2013 Jul;49(4):764–6.

32. Cohen J. Statistical power analysis for the behavioral sciences. 2nd ed. Hillsdale, N.J: L. Erlbaum Associates; 1988. 567 p.

33. Vaishya R, Misra A, Vaish A, Ursino N, D’Ambrosi R. Hand grip strength as a proposed new vital sign of health: a narrative review of evidences. J Health Popul Nutr. 2024 Jan 9;43(1):7.

34. Hackney KJ, Ploutz-Snyder LL. Unilateral lower limb suspension: integrative physiological knowledge from the past 20 years (1991–2011). Eur J Appl Physiol. 2012 Jan 1;112(1):9–22.

35. Deschenes MR, Holdren AN, McCoy RW. Adaptations to short-term muscle unloading in young and aged men. Med Sci Sports Exerc. 2008 May;40(5):856–63.

36. Hvid L, Aagaard P, Justesen L, Bayer ML, Andersen JL, Ørtenblad N, et al. Effects of aging on muscle mechanical function and muscle fiber morphology during short-term immobilization and subsequent retraining. J Appl Physiol (1985). 2010 Dec;109(6):1628–34.

37. O’Bryan SJ, Critchlow A, Fuchs CJ, Hiam D, Lamon S. The contribution of age and sex hormones to female neuromuscular function across the adult lifespan. The Journal of Physiology. 2025 May 11;JP287496.

38. Preobrazenski N, Janssen I, McGlory C. The effects of single-leg disuse on skeletal muscle strength and size in the nonimmobilized leg of uninjured adults: a meta-analysis. J Appl Physiol (1985). 2023 Jun 1;134(6):1359–63.

39. Carr JC, Voskuil CC, Andrushko JW, MacLennan RJ, DeFreitas JM, Stock MS, et al. Cross-education attenuates muscle weakness and facilitates strength recovery after orthopedic immobilization in females: A pilot study. Physiol Rep. 2025 Apr;13(8):e70329.

40. Wegrzynowicz AK, Eyvazzadeh A, Beckley A. Current Ovulation and Luteal Phase Tracking Methods and Technologies for Fertility and Family Planning: A Review. Semin Reprod Med. 2024 Jun;42(2):100–11.

41. Elliott-Sale KJ, Altini M, Doyle-Baker P, Ferrer E, Flood TR, Harris R, et al. Why We Must Stop Assuming and Estimating Menstrual Cycle Phases in Laboratory and Field-Based Sport Related Research. Sports Med. 2025 Mar 14;

42. Logeson ZS, MacLennan RJ, Abad GKB, Methven JM, Gradl MR, Pinto MD, et al. The impact of skeletal muscle disuse on distinct echo intensity bands: A retrospective analysis. PLoS One. 2022;17(1):e0262553.

